# Genome Sequence and Annotation the B3 Mycobacteriophage Phayeta

**DOI:** 10.1101/2023.07.21.550035

**Authors:** Emily Bishop, Warren Earlely, Alexandra Greco, Emma Hofseth, Emma Kinerson, Brandon Lafayette, Nestor Llanot-Arocho, Brittney Mazen, Megan Cevasco, Daniel C. Williams

**Affiliations:** Department of Biology, Coastal Carolina University, Conway, SC

## Abstract

Phayeta is a siphovirus extracted from soil near Myrtle Beach, South Carolina which infects *Mycobacterium smegmatis*. Annotation of the 68,700 base-pair circularly permuted genome identified 104 predicted protein-encoding genes, 36 of which have functional assignments.

---

*Mycobacterium smegmatis* is a genetically tractable model of *M. tuberculosis* and phages that infect *M. smegmatis* have potential to be therapeutically useful (1, 2). Mycobacteriophage Phayeta was extracted from a soil sample taken near Myrtle Beach, South Carolina (Global Positioning Coordinates 33.75493N, 78.90047 W), using standard protocols (3). Briefly, the soil sample was washed in 7H9 liquid media, the wash filtered (0.22 um), and the filtrate mixed with *M. smegmatis* for 15 minutes before being plated in top agar and incubated at 37°C for 48 hours. Phayeta produced small clear plaques that are characteristic of non-temperate phages (Figure 1a). Plaque size was measured with <u>ImageJ</u> and ranged from 0.5 mm to 1.7 mm with an average of 1.0 ± 0.2 mm (mean ± standard deviation, n=50) (4, 5). Negative-staining transmission electron microscopy of Phayeta (Figure 1b) revealed a siphovirus morphology consisting of an isometric capsid (diameter = 76.6 ± 3.6 nm) and long flexible tail (274.7 ± 11.6, n = 9 viral particles).

**Figure 1.**
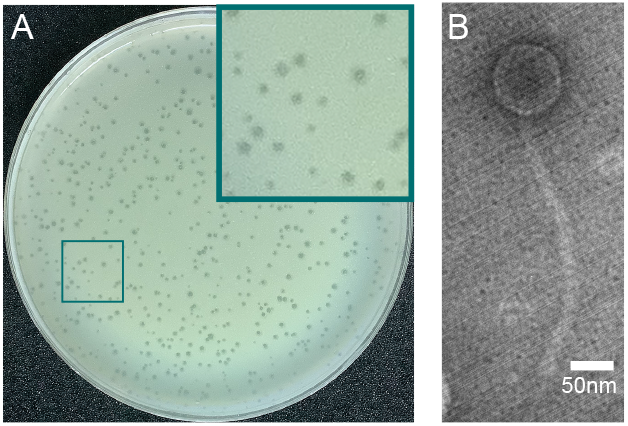
The mycobacteriophage Phayeta. (A) Plaque assay. Purified samples of Phayeta were mixed with *M. semgmatis* MC^2^155, then combined with molten 7H9 top-agar and poured on 90 mm plates. Incubation at 37°C for 48 hours resulted in the production of clear small plaques. (B) Transmission electron microscopy of Phayeta negatively stained with 1 % uranyl acetate and imaged using a JEOL JEM 1230 at 100 kV.

Phayeta DNA was extracted from high-titer lysates using the Wizard DNA Clean-Up Kit (Promega) and analyzed by agarose gel electrophoresis. Intact DNA was prepared with the Ultra II Library Kit (New England Biolabs) and sequenced on the Illumina MiSeq platform (v3 reagents) at the Pittsburgh Bacteriophage Institute. A total of 446,915 single-end 150 base reads provided 278x coverage of the Phayeta genome. Sequence reads were assembled and verified using Newbler (v2.9) and Consed (v29) (6). The genome is 68,700 base-pairs long, has 67.5% GC content, and contains circularly permuted genomic ends. Comparison of gene content similarity of phages within the Actinobacteriophage Database, (phagesdb.org) placed Phayeta in subcluster B3 (7). Among this cluster, Phayeta shares the greatest gene content similarity with Casbah (94.6%) and Kronus (91.7%).

Initial draft annotation of Phayeta was done using the GenMark (v2.5), Glimmer (v3.02), and ARAGORN algorithms within DNA Master (v5.23.56) (8–10). Subsequent manual annotations employed Phamerator (v7), <u>Starterator</u> (v1.3), and <u>PECAAN</u>. (11). Both PhagesDB and NCBI nonredundant databases were searched for sequence-based homology comparison via BLAST (7, 12). Predicted protein structure comparisons used HHPRED (v3.18) to search the PDB mmCIF70, Pfam-A, and NCBI Conserved Domain databases using default parameters. A total of 104 protein encoding genes were identified, 36 of which have a functional assignment.

Most genes on the left arm of Phayeta encode structural or assembly proteins with the right arm containing genes involved in DNA replication. Consistent with a non-temperate lifestyle, Phayeta lacks genes encoding an identifiable integrase or repressor. Both a Lysin A and Lysin B are present. Although homology searching failed to identify a holin encoding gene, a small cluster of transmembrane-segment containing genes (gp17-gp20) have sequence and syntenic similarities with genes of *Gordonia* phages that are a predicted holin cassette (13). There are five minor tail proteins, including gp33 which shows full-length similarity to Corofin gp33. Both these phages have a single nucleotide indel polymorphism in this gene, but at different locations suggesting distinct mutational events.

## Nucleotide sequence accession numbers

Phayeta is available at GenBank with Accession No. <u>OR159655</u> and Sequence Read Archive No. <u>SRX20630268</u>

## Acknowledgements

We thank Dan Russell and Ching-Chung Ko for sequencing and genome assembly, Julian Smith III and Victoria Frost of Winthrop University for their electron microscopy service, and the Howard Hughes Medical Institute SEA-PHAGES program and Coastal Carolina University Biology Department for support.

